# Targeting Neutrophils Extracellular Traps (NETs) reduces multiple organ injury in a COVID-19 mouse model

**DOI:** 10.1101/2022.04.27.489676

**Authors:** Flavio P. Veras, Giovanni F. Gomes, Bruna M. S. Silva, Cicero J. L. R. Almeida, Camila Meirelles S. Silva, Ayda H. Schneider, Emily S. Corneo, Caio S. Bonilha, Sabrina S. Batah, Ronaldo Martins, Eurico Arruda, Alexandre T. Fabro, José C. Alves-Filho, Thiago M. Cunha, Fernando Q. Cunha

## Abstract

COVID-19 is characterized by severe acute lung injury, which is associated with neutrophils infiltration and release of neutrophil extracellular traps (NETs). COVID-19 treatment options are scarce. Previous work has shown an increase in NETs release in the lung and plasma of COVID-19 patients suggesting that drugs that prevent NETs formation or release could be potential therapeutic approaches for COVID-19 treatment. Here, we report the efficacy of NET-degrading DNase I treatment in a murine model of COVID-19. DNase I decreased detectable levels of NETs, improved clinical disease, and reduced lung, heart, and kidney injuries in SARS-CoV-2-infected K18-hACE2 mice. Furthermore, our findings indicate a potential deleterious role for NETs lung tissue *in vivo* and lung epithelial (A549) cells *in vitro*, which might explain part of the pathophysiology of severe COVID-19. This deleterious effect was diminished by the treatment with DNase I. Together, our results support the role of NETs in COVID-19 immunopathology and highlight NETs disruption pharmacological approaches as a potential strategy to ameliorate COVID-19 clinical outcomes.

## INTRODUCTION

Coronavirus diseases 2019 (COVID-19) is an infection caused by the severe acute respiratory syndrome coronavirus 2 (SARS-CoV-2) (1, 2) and pulmonary-related symptoms are one of its hallmarks (3, 4). Neutrophils have been described as indicators of severity of respiratory symptoms and poor COVID-19 prognosis (5–7). Neutrophil extracellular traps (NETs) are one of the most relevant effector mechanisms of neutrophils in inflammatory diseases, playing central role in organs damage (8–10).

NETs are web-like structures of extracellular DNA fibers containing histones and granule-derived enzymes, such as myeloperoxidase (MPO), neutrophil elastase, and cathepsin G (11, 12). The formation of NETs is known as NETosis, and it starts with neutrophil activation by pattern recognition receptors (TLRs, e.g) or chemokines. The process is followed by ROS production and calcium mobilization, which leads to the activation of protein arginine deiminase 4 (PAD-4) (13). The activation of the neutrophil elastase also plays a role in NETs production in inflammatory responses (14–17).

Elevated levels of NETs are found in the blood, thrombi, and lungs of patients with severe COVID-19, suggesting that neutrophils and NETs may play an important role in the pathophysiology of COVID-19 (8–10).

Drug repurposing is a key strategy to accelerate discovery of new effective treatments for COVID-19 (18). In this context, as the literature shows evidence of the role of NETs in COVID-19, DNase I, an FDA-approved drug that degrades NETs could be proposed as a potential new candidate for COVID-19 treatment (19–21).

Here, we demonstrate that DNase I treatment decreases the concentration of NETs in the plasma and lungs of SARS-CoV-2-infected mice and ameliorates experimental COVID-19. These findings highlight the importance of NETs inhibitors as a potential therapeutic approach for COVID-19 treatment.

## RESULTS

### DNase I reduces clinical outcomes in an experimental model of COVID-19

To investigate the role of NETs in COVID-19 pathophysiology we used K18-hACE2 mouse intranasally infected with SARS-CoV-2; (22, 23). Infected mice were treated with saline or DNAse I (10 mg/kg; s.c.) administered 1 h before virus infection and once daily up to 4 days after infection (**Figure 1A)**. We evaluated body weight loss from baseline measure (basal) and clinical scores, as read out of disease progression. Our data showed that DNase I treatment attenuates both body weight loss and clinical score caused by SARS-CoV-2 infection compared to vehicle-treated mice (**Figure 1, B and C**).

**Figure 1.**
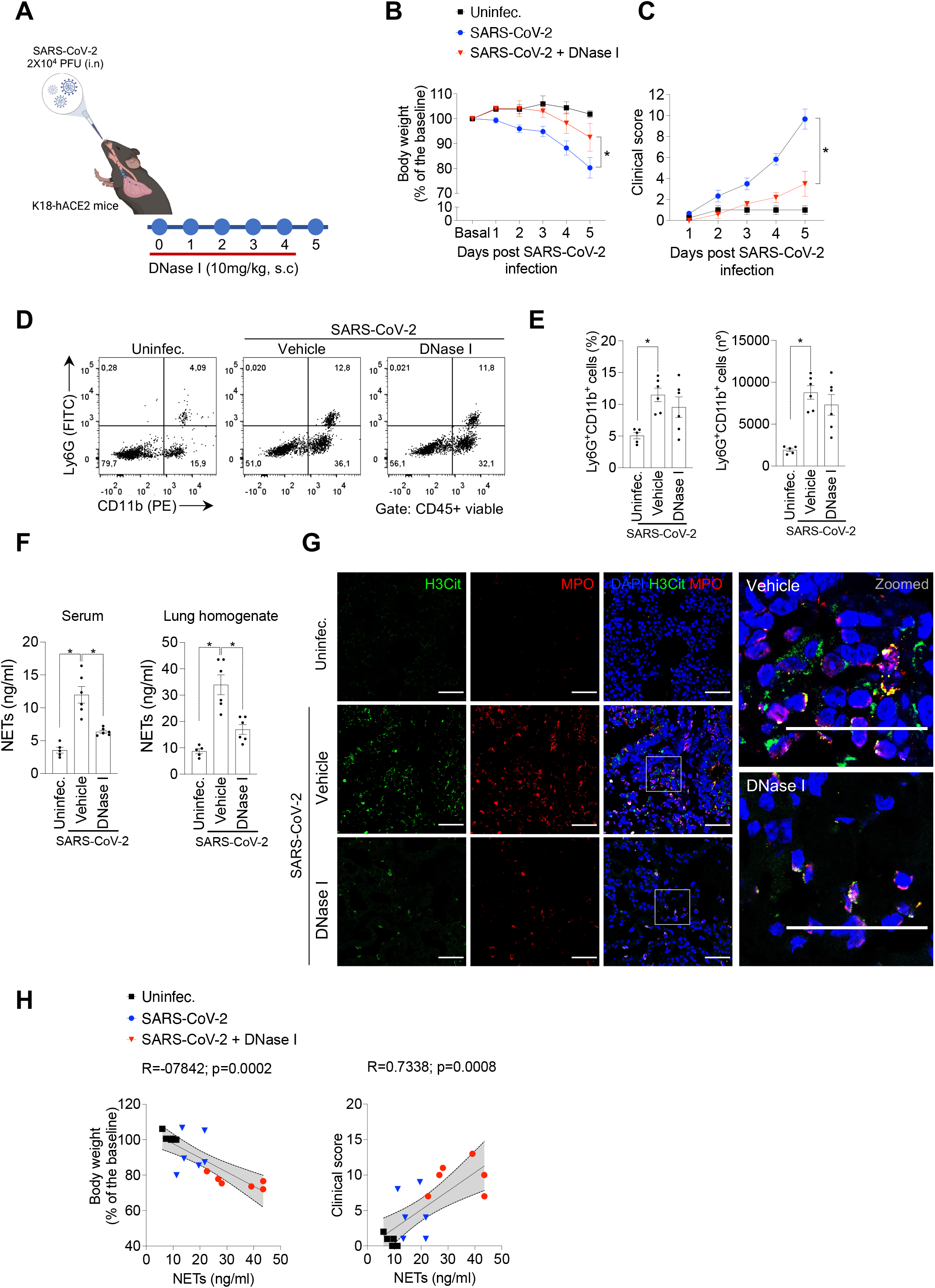
DNase I reduces clinical outcomes in a COVID-19 experimental model. **(a)** K18-hACE2 mice (n=6) were intranasally (i.n) inoculated with SARS-CoV-2 (2×10^4^ PFU) and treated with DNase I (10mg/kg, s.c) for 5 days. Created with Biorender.com. Uninfected group was used as negative control (n=5). Weight loss and clinical scores were analyzed **(b, c)**. Representative dot plots **(d)**, frequency and total number **(e)** of Ly6G+CD11b+ cells assessed by flow cytometry. **(f)** NETs quantification by MPO-DNA PicoGreen assay from serum and lung tissue homogenate. **(g)** Immunofluorescence analysis of H3Cit (green) and MPO (red) expression from the lung of SARS-CoV-2-infected mice and treated with DNase I. DAPI (blue) was used for nuclei staining. Scale bar indicates 50 μm. **(h)** Pearson’s correlation between body weight or clinical score and NETs concentration in lung homogenate. Data are representative of at least two independent experiments and are shown as mean ± SEM. P values were determined by two-way **(b, c)** and one-way ANOVA **(e, f)**.

We observed an increase in NETs concentration **(Figure 1, F and G)** and in the number of neutrophils in the lung of infected mice upon induction of the COVID-19 model **(Figure 1, D and E)**. These data corroborate some findings published by either our group and other authors, showing the presence of systemic levels of NETs, as well as localized in pulmonary tissue from COVID-19 patients (8, 9, 24).

DNase I treatment did not alter the number of neutrophils that infiltrated into lung tissue of SARS-CoV-2 infected mice. The FACS analysis showed that the frequency and absolute numbers of neutrophils (Ly6G+CD11b+ cells; **Figure 1, D and E**) were not altered by the treatment. Similar results were observed when analyzing CD45+ cell populations (**Supplemental Figure 1**). However, we demonstrate that DNAse I treatment of SARS-CoV-2-infected mice resulted in a substantial reduction of NETs production that associates with lower disease score (**Figure 1H**). These data suggest that NETs play a crucial role in SARS-CoV-2 infection.

### Inhibition of NETs ameliorates lung pathology in SARS-CoV-2-infected mice

Lung inflammation is the primary cause of life-threatening respiratory disorder in critical and severe forms of COVID-19 (3, 4, 25). NETs have been identified in the lung tissue of SARS-CoV-2 infected patients and lung injury experimental animal models (8, 9, 26). Thus, we next investigated the effect of NETs degradation by DNase I treatment of the COVID-19 mouse model.

We noticed an extensive injury in lung tissue of K18-hACE2 infected mice with interstitial leukocyte infiltration. The alveolar units showed architectural distortion compromising the alveolar capillary barrier. DNase I treatment was able to prevent lung damage caused by SARS-CoV-2 infection (**Figure 2A**). Moreover, analyzing the area fraction, as score of septal thickness, DNase I-treatment reduced the area fraction in SARS-CoV-2-infected mice and was associated with lower concentration of NETs released **(Figure 2B).**

**Figure 2.**
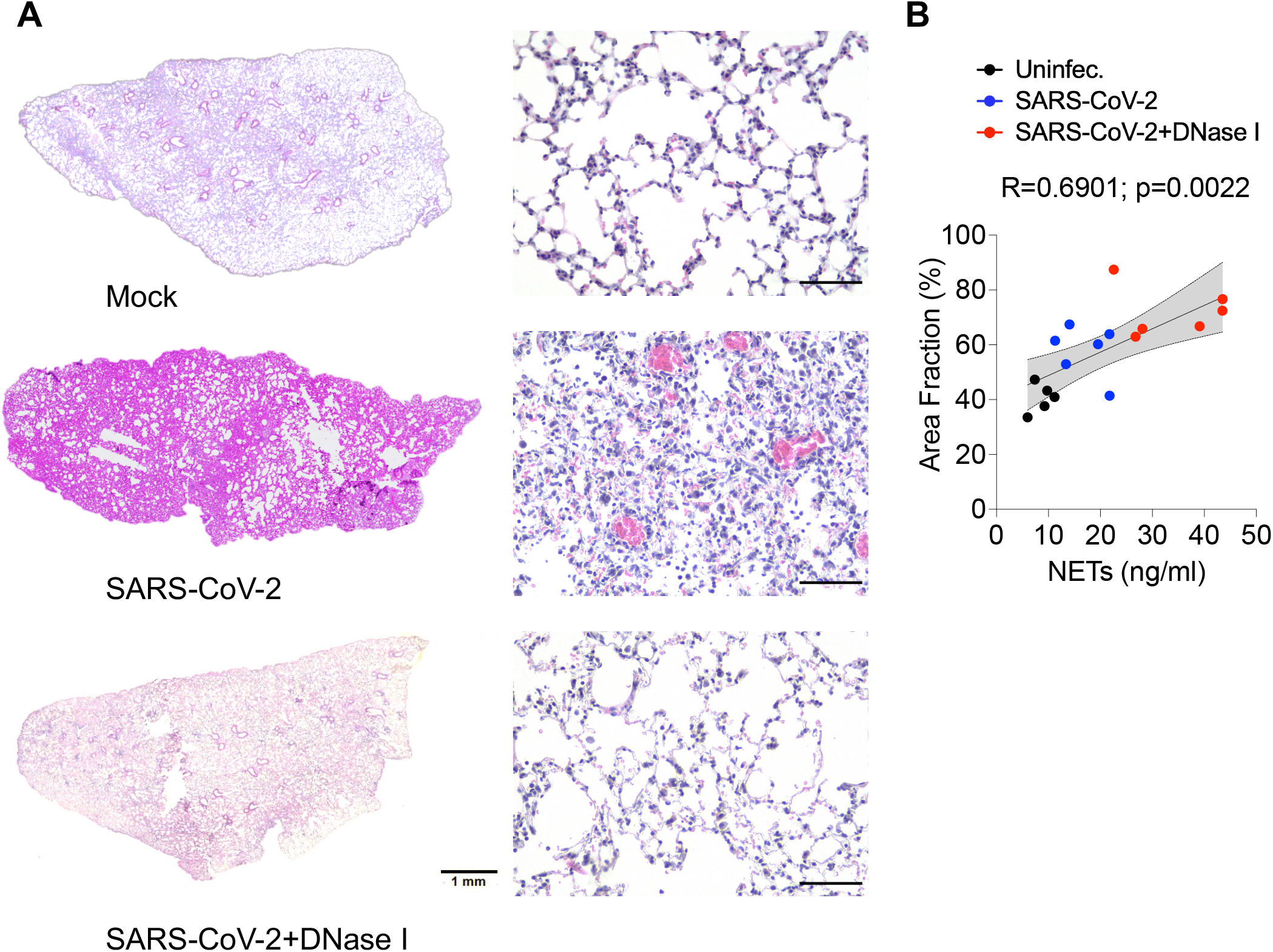
Pharmacological degradation of NETs ameliorates lung pathology upon SARS-CoV-2 infection *in vivo.* K18-hACE2 mice (n=6) were intranasally (i.n) inoculated with SARS-CoV-2 (2×10^4^ PFU) and treated with DNase I (10mg/kg, s.c) for 5 days. **(a)** Representative H&E staining from the lung of mock or SARS-CoV-2-infected mice and treated with DNase I. **(b)** Pearson correlation between the area fraction and NETs concentration in lung homogenate. Data are representative of two independent experiments and are shown as mean ± SEM. P values were determined by one-way ANOVA.

### DNase I attenuates extrapulmonary injuries in the COVID-19 mouse model

COVID-19 is a systemic viral disease that can affect other vital organs besides the lungs, such as heart and kidney (27–30). To investigate this context, we harvested heart and kidney of animals 5 days post infection and evaluated whether NETs inhibition could change this phenotype. We observed that heart of mice infected with SARS-CoV-2 showed pathological changes in cardiac tissue with diffuse and sparse cardiac inflammatory cell infiltration and perivascular injury (**Figure 3A**). In addition, plasma Creatine Kinase MB (CK-MB) concentration in the infected group was higher than in control mice (**Figure 3B**). Interestingly, DNase I treatment attenuated the pathology and CK-MB concentration.

**Figure 3.**
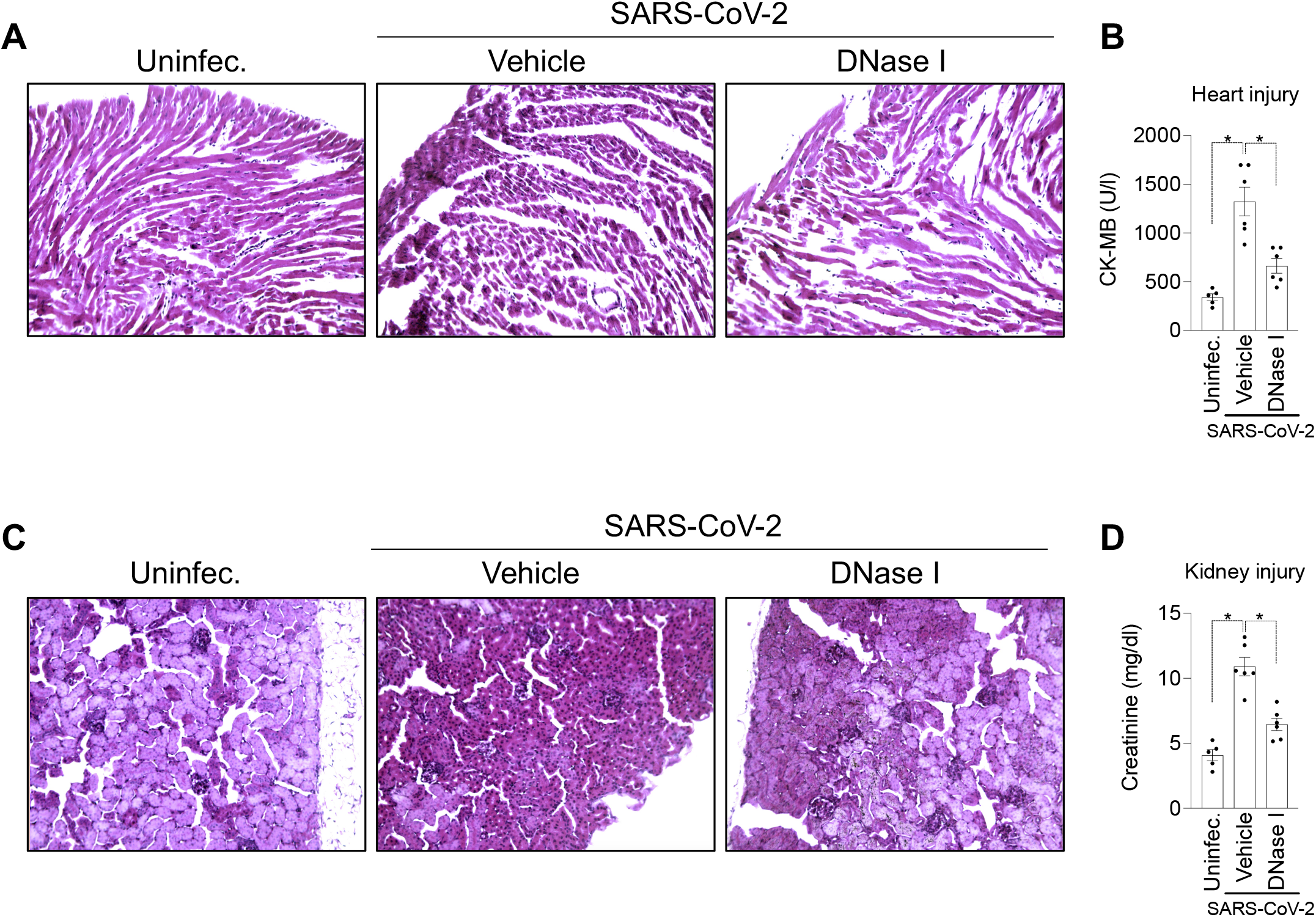
NETs degradation attenuates heart and kidney failure in a COVID-19 experimental model. K18-hACE2 mice (n=6) were intranasally (i.n) inoculated with SARS-CoV-2 (2×10^4^ PFU) and treated with DNase I (10mg/kg, s.c) for 5 days. Representative H&E staining from heart **(a)** and kidney **(c)** of mock, SARS-CoV-2-infected mice and treated with DNase I. Serum levels of creatine kinase-MB (CK-MB) **(b)** and creatinine **(d)** determined by colorimetric assays. Data are representative of two independent experiments and are shown as mean ± SEM. P values were determined by one-way ANOVA.

Finally, we aimed to investigate whether SARS-CoV-2 infection induces kidney damage in K18-hACE2 mice. Consistent with lung and heart pathological changes, we found the presence of ischemic tubulointerstitial nephritis in the COVID-19 model, mimicking acute tubular necrosis with cellular glomerulitis (**Figure 3C)**. Furthermore, the levels of creatinine in the blood of infected mice were higher in comparison with uninfected animals (**Figure 3D)**. As expected, DNase I treatment reduced kidney injury (**Figure 3, C and D)**. Collectively, these findings indicate that pharmacological inhibition of NETs with DNase I prevents multi-organ dysfunction in COVID-19.

### DNase I prevents NETs-induced apoptosis of lung tissue in SARS-CoV-2-infected mice

To understand the possible role of NETs in the pathophysiology of COVID-19, we explored the hypothesis that NETs could be involved in the lung damaged accompanied by the disease, as previously described (9, 31).

To this end, lung analysis of k18-hACE2 mice infected with SARS-CoV-2 reveals a massive presence of apoptotic cells after 5 days of infection. DNase I treatment was able to prevent lung apoptosis caused by COVID-19 mouse model (**Figure 4A**). Our analysis shown a reduction of approximately 60% in TUNEL positive cells after NETs degradation (**Figure 4B)**.

**Figure 4.**
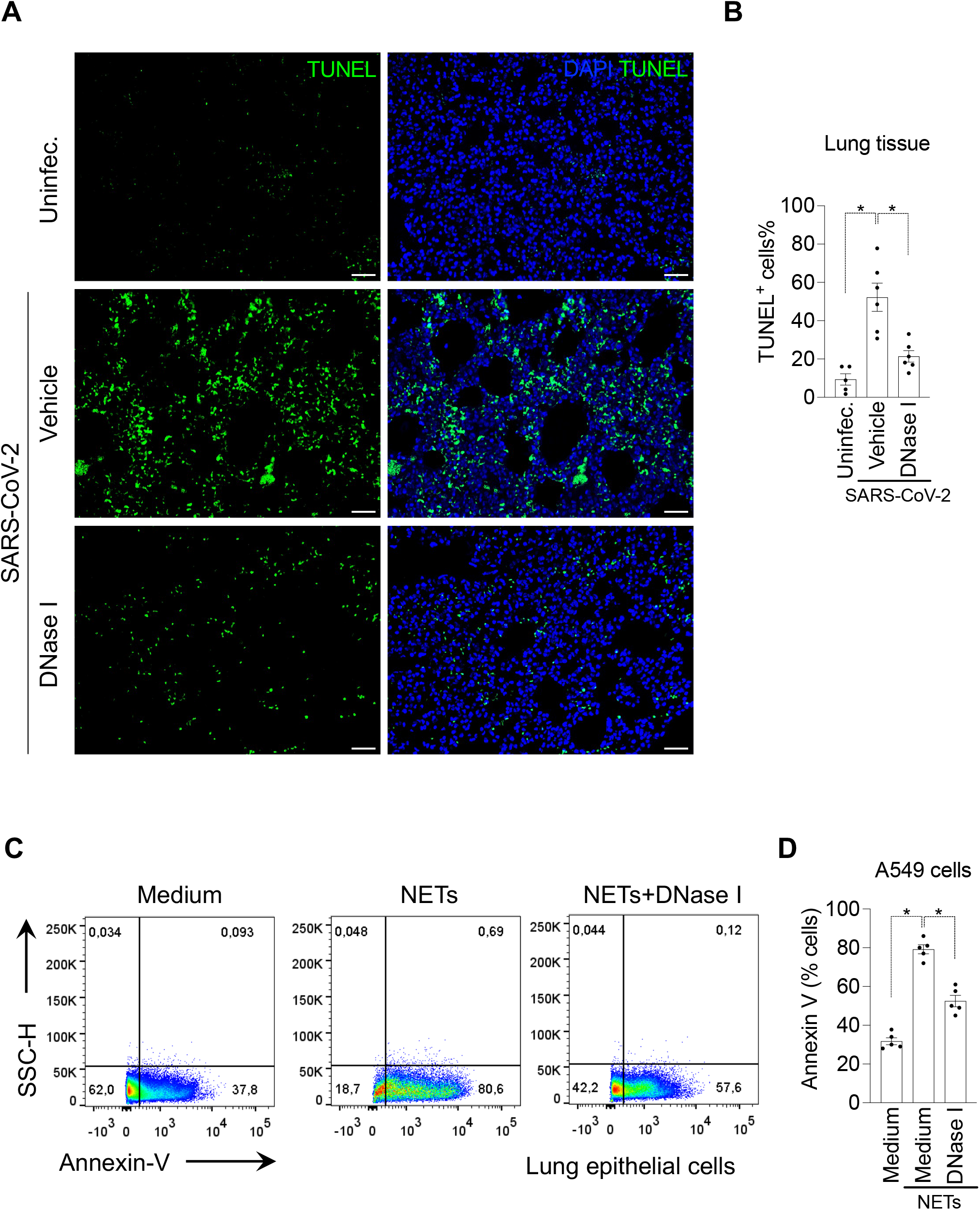
DNase I prevents apoptosis of lung tissue and lung epithelial cells. K18-hACE2 mice (n=6) were intranasally (i.n) inoculated with SARS-CoV-2 (2×10^4^ PFU) and treated with DNase I (10mg/kg, s.c) for 5 days. (**a**) TUNEL staining (green) for detection of apoptotic cells *in situ* from lung tissue of mice. (**b**) Percentage of TUNEL positive cells in lung tissue. NETs were purified from healthy neutrophils and stimulated with PMA (50 nM) for 4 h at 37°C **(c)** Representative dot plots of FACS analysis of Annexin V+ A549 cells incubated with purified NETs (10 ng/ml) pretreated, or not, with DNase I (0.5 mg/ml) for 24 h at 37°C. **(d)** Frequency of Annexin V+ A549. Data are representative of two independent experiments and are shown as mean ± SEM. P values were determined by one-way ANOVA followed by Bonferroni’s post hoc test.

In our next step, purified NETs from healthy human neutrophils were incubated with A549 cells, a human alveolar basal epithelial cell, and cell viability was determined (annexin V+ cells) by flow cytometry.

We found that exposure of A549 cells to purified NETs significantly increased the percentage of apoptotic cells in comparison with untreated cells. Importantly, the pretreatment of purified NETs with DNase I prevented NETs-induced epithelial **(Figure 4, C and D)** apoptosis to the similar levels observed in untreated cells. Taken together, these results are evidence that DNase I exhibits a protective effect in NETs deleterious functions in lung tissue and could be a potential strategy to block organ injury during COVID-19.

## DISCUSSION

While the number of patients with COVID-19 is growing worldwide, there is no effective treatment for the disease (32, 33). Thus, the understanding of the mechanisms by which the hosts deal with SARS-CoV-2 virus could allow the development of new therapeutic strategies aiming to prevent tissue injuries triggered by the infection. Here, we report that in a COVID-19 mouse model, NETs are released systemically and in higher concentration in the lungs of K18-hACE2 mice. Moreover, DNase I treatment reduced multi-organ lesions and improved outcomes associated NETs released.

The increase in the number of circulating neutrophils is an indicator of a worse outcome of COVID-19 (34). In 2004, Brinkmann et al. described, for the first time, that NETs are released by neutrophils, and work as a microbicidal mediator (11). However, following studies demonstrated that NETs have a dual biological role. Besides their microbicidal ability, NETs mediate lesions observed in several inflammatory diseases, including rheumatoid arthritis, lupus, diabetes, and sepsis (8, 10, 35–39). In fact, inhibition of NETs production prevented lung, heart and liver lesions observed in experimental sepsis (38–40).

There are several stimuli that trigger NETs release, including pathogen-associated molecular patterns (PAMPs), damage-associated molecular patterns (DAMPs) and inflammatory mediators, such as cytokines and chemokines (41–43). Our group have previously demonstrated that SARS-CoV-2 can directly infect human neutrophils and is key to trigger NETs production. The first step in neutrophil infection by SARS-CoV-2 is the interaction of the virus with ACE2 and TMPRSS2 expressed in the surface of human neutrophils (8). It is possible to speculate the participation of the cytokines and chemokines released by host cells as activators (44, 45) of NETs production by the neutrophils in the K18-hACE2 model; however, this deserves future investigation.

The literature describes that NETs can present direct cytotoxic effects on different mammalian cell types, including epithelial and endothelial cells, inducing apoptosis or necrosis (12, 46). Moreover, NETs could also activate different PRR receptors, such as toll-like receptor (TLR)-4 and 9, which mediate the release of inflammatory mediators; in turn, amplifying the direct effects of NETs (35). In this context, during COVID-19, apoptosis of lung epithelial was previously observed. These events are capable of compromising the lung function, worsening the severity of the disease (3, 4). Considering these findings, in the present study, we observed that DNase I prevented apoptosis in lung tissue from SARS-CoV-2-infected mice. In accordance observed in COVID-19 model, isolated NETs from culture of PMA-stimulated human neutrophils induced *in vitro* apoptosis of epithelial cells with reversal homeostasis in presence of DNase I. Extending this finding to COVID-19 disease, it is possible to suggest that the reduction of viability of the lung cells is a consequence of local production of NETs. In this line, we observed the presence of neutrophils releasing NETs, as well as high concentration of NETs in the lung of SARS-CoV-2-infected mice. Finally, supporting these finds, we observed the presence of neutrophils releasing NETs in the lung tissue of infected mice, detected in the alveolar space. Disease progression was prevented with DNase I treatment *in vivo.* This observation is tightly correlated with the development of lung injury, suggesting that strategies to reduce NETs levels could have favorable effects on recovering lung function. Together, our findings demonstrate the potentially deleterious role of NETs during COVID-19 and support the use of inhibitors of NETs, such as DNase I as a strategy to ameliorate multi-organ damage during COVID-19.

## MATERIALS AND METHODS

### K18-hACE2 mice

K18-hACE2 humanized mice (B6.Cg-Tg(K18-ACE2)2Prlmn/J) were obtained from The Jackson Laboratory and were bred in the *Centro de Criação de Animais Especiais* (Ribeirão Preto Medical School/University of São Paulo). This mouse strain has been previously used as model for SARS-CoV-2-induced disease and it presents signs of diseases, and biochemical and lung pathological changes compatible with the human disease (23). Mice had access to water and food ad libitum. The manipulation of these animals was performed in Biosafety Levels 3 (BSL3) facility and the study was approved by Ethics Committee on the Use of Animals of the Ribeirão Preto Medical School, University of São Paulo (#066/2020).

### DNase I treatment in SARS-CoV-2 experimental infection

Male K18-hACE2 mice, aged 8 weeks, were infected with 2×10^4^ PFU of SARS-CoV-2 (in 40 μL) by intranasal route. Uninfected mice (n=5) were given an equal volume of PBS through the same route. On the day of infection, 1 h before virus inoculation, animals were treated with DNase I (10 mg/kg, s.c., Pulmozyme, Roche) (n = 6) or vehicle (PBS, s.c.) (n = 6). DNase I was also given once a day until 5 days post-infection. Body weight was evaluated on the baseline and on all days post-infection. The right lung was collected, harvested, and homogenized in PBS with steel glass beads. The homogenate was added to TRIzol reagent (1:1), for posterior viral titration via RT-qPCR, or to lysis buffer (1:1), for ELISA assay, and stored at −70 °C. The left lung was collected in paraformaldehyde (PFA, 4%) for posterior histological assessment.

### H&E staining and lung pathology evaluation

Five μm lung, heart and kidney slices were submitted to Hematoxylin and Eosin staining. A total of 10 photomicrographs in 40X magnification per animal were randomly obtained using a microscope ScanScope (Olympus) and Leica. Morphometric analysis was performed in accordance with the protocol established by the American Thoracic Society and European Thoracic Society (ATS/ERS) (47).

### NETs quantification

Plasma or homogenate from lung were incubated overnight in a plate pre-coated with anti-MPO antibody (Thermo Fisher Scientific; cat. PA5-16672) at 4°C. The plate was washed with PBS-T (Phophate-Buffered Saline with Tween 20). Next, samples were incubated overnight at 4°C. Finally, the plate with samples was washed and over MPO-bound DNA was quantified using the Quant-iT PicoGreen kit (Invitrogen; cat. P11496).

### Immunofluorescence and confocal microscopy

Lungs were harvested and fixed with PFA 4%. After dehydration and paraffin embedding, 5 μm sections were prepared. The slides were deparaffinized and rehydrated by immersing the through Xylene and 100% Ethanol 90% for 15 minutes, each solution. Antigen retrieval was performed with 1.0 mM EDTA, 10 mM Trizma-base, pH 9.0 at 95°C for 30 minutes. Later, endogenous peroxidase activity was quenched by incubation of the slides in 5% H2O2 in methanol for 15 minutes. After blocking with IHC Select Blocking Reagent (Millipore, cat. 20773-M) for 2 hours at room temperature (RT), the following primary antibodies were incubated overnight at 4°C: rabbit polyclonal anti-myeloperoxidase (anti-MPO; Thermo Fisher Scientific; cat. PA5-16672; 1:100) and rabbit polyclonal, anti-histone H3 (H3Cit; Abcam; cat. ab5103; 1:100). The slides were then washed with TBS-T (Tris-Buffered Saline with Tween 20) and incubated with secondary antibodies alpaca anti-rabbit IgG AlexaFluor 488 (Jackson ImmunoReseacher; Cat. 615-545-215; 1:1000) and alpaca anti-rabbit IgG AlexaFluor 594 (Jackson ImmunoReseacher; Cat. 611-585-215; 1:1000). Autofluorescence was quenched using the TrueVIEW Autofluorescence Quenching Kit (Vector Laboratories, cat. SP-8400-15). Slides were then mounted using Vectashield Antifade Mounting Medium with DAPI (Vector Laboratories, Cat# H-1200-10). Images were acquired by Axio Observer combined with a LSM 780 confocal microscope (Carl Zeiss) at 63X magnification at the same setup (zoom, laser rate) and tile-scanned at 4 fields/image. Images were analyzed with Fiji by Image J.

### Measurement of organ damage biomarkers

Renal dysfunction was assessed by the levels of blood creatinine; and creatine kinase-MB was used as an index of cardiac lesions. The determinations were performed using a commercial kit (Bioclin).

### Neutrophils isolation and NETs purification

Peripheral blood samples were collected from healthy controls by venipuncture and the neutrophil population was isolated by Percoll density gradient (GE Healthcare; cat. 17-5445-01). Isolated neutrophils (1.5 × 10^7^ cells) were stimulated with 50 nM of PMA (Sigma-Aldrich; cat. P8139) for 4 h at 37°C. The medium containing NETs was centrifuged at 450 g to remove cellular debris for 10 minutes, and NETs-containing supernatants were collected and centrifuged at 18,000 g for 20 minutes. Supernatants were removed, and DNA pellets were resuspended in PBS. NETs were then quantified with a GeneQuant (Amersham Biosciences Corporation).

### Apoptosis assay

Lung tissue were harvested for detection of apoptotic cells *in situ* with Click-iT Plus TUNEL Assay Alexa Fluor 488, according to the manufacturer’s instructions (Thermo Fisher Scientific; cat. C10617). Human alveolar basal epithelial A549 cells (5×10^4^) were maintained in DMEM and cultured with purified NETs (10 ng/ml) pretreated, or not, with DNase I (0.5 mg/ml; Pulmozyme, Roche). The cultures were then incubated for 24 h at 37°C. Viability was determined by flow cytometric analysis of Annexin V staining.

### Flow cytometry

Lung tissue was harvested and digested with type 2 collagenase to acquire cell suspensions. Cells were then stained with Fixable Viability Dye eFluor 780 (eBioscience; cat. 65–0865-14; 1:3,000) and monoclonal antibodies specific for CD45 (BioLegend; clone 30-F11; cat. 103138; 1:200), CD11b (BD Biosciences; clone M1/70; cat. 553311) and Ly6G (Biolegend; clone 1A8; cat. 127606) for 30 min at 4°C. A549 cells (1×10^5^) were stained with FITC ApoScreen Annexin V Apoptosis Kit (SouthernBiotech; cat. 10010–02), according to the manufacturer’s instructions. Data were collected on a FACSVerse (BD Biosciences) and analyzed with FlowJo (TreeStar).

### Statistics

Statistical significance was determined by either two-tailed unpaired Student t test, one-way or two-way ANOVA followed by Bonferroni’s post hoc test. P<0.05 was considered statistically significant. Statistical analyses and graph plots were performed and built with GraphPad Prism 9.3.1 software.

## Acknowledgments

We are grateful to Marcella Daruge Grando, Roberta Rosales, Soraya Jabur Badra, Andreia Nogueira, and Juliana Trench Abumansur for technical assistance. This research was supported by Fundação de Amparo à Pesquisa do Estado de São Paulo (FAPESP) grants (2013/08216-2 and 2020/05601-6), Conselho Nacional de Desenvolvimento Científico e Tecnológico (CNPq) and Coordenação de Aperfeiçoamento de Pessoal de Nível Superior (CAPES) grants.

## Author contributions

F.P. Veras, G.F. Gomes, F.Q. Cunha, T.M. Cunha, J.C. Alves-Filho, E. Arruda and A.T. Fabro contributed to the study design. F.P. Veras, G.F. Gomes, E.S. Corneo, R. Martins and, B.M.S. Silva performed SARS-CoV-2 K18-hACE2 mice infection *in vivo.* F.P. Veras performed immunostaining and confocal analysis. A.H. Schneider, C.J.L.R. Almeida, and C.M.S. Silva performed NETs quantification. S.S. Batah, A.T. Fabro contributed to histopathological analysis. F.P. Veras and B.M.S. Silva performed A549 epithelial cell damage assay and FACS analysis. A.H. Schneider performed the purification of NETs. F.P. Veras, R. Martins, C.S Bonilha, E. Arruda F.Q. Cunha, T.M. Cunha, and J.C. Alves-Filho, wrote the manuscript. All authors read and approved the manuscript.

---

**Figure S1.**
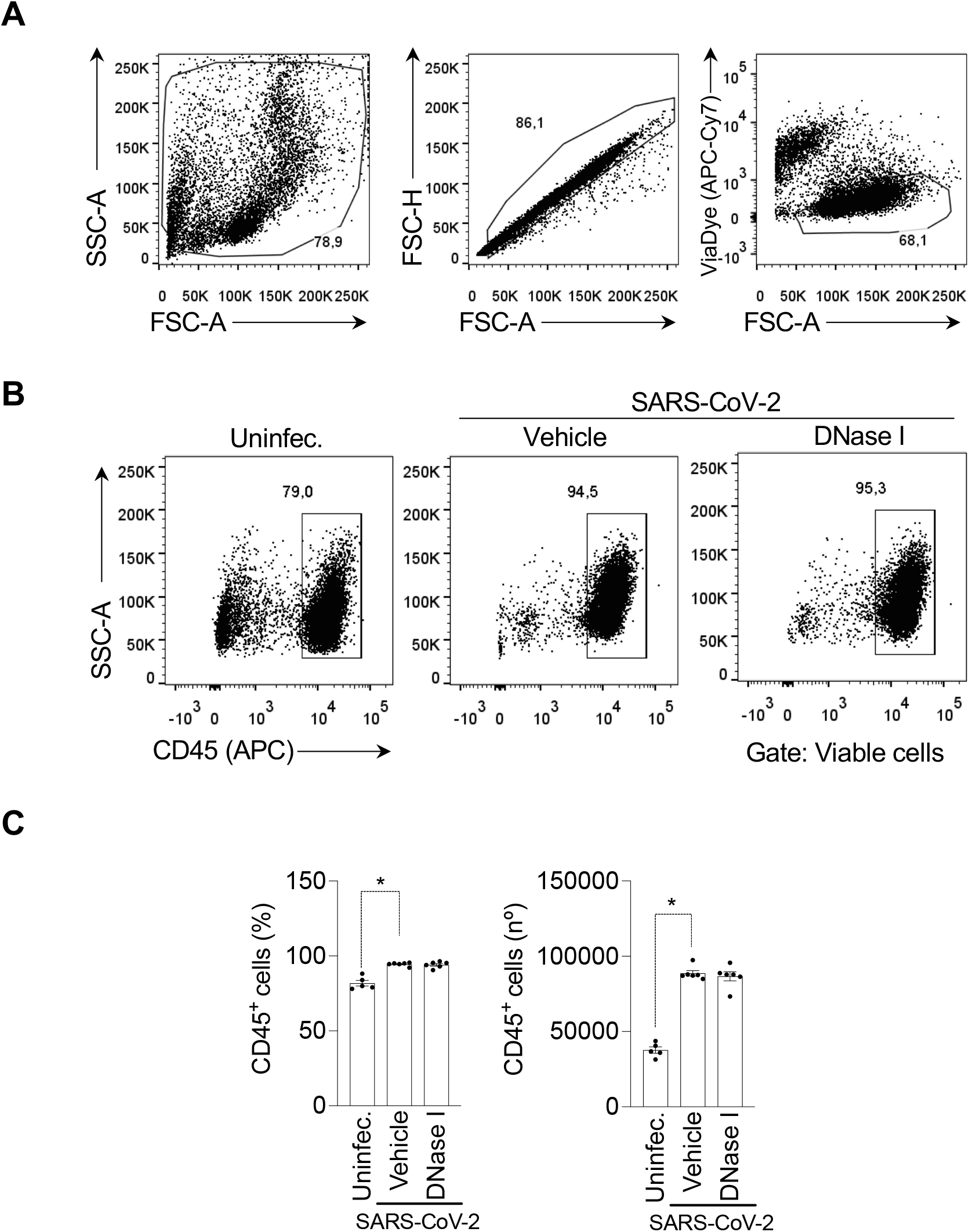
DNase I treatment does not alter leukocyte accumulation in the lungs of SARS-CoV-2-infected mice. K18-hACE2 mice (n=6) were intranasally (i.n) inoculated with SARS-CoV-2 (2×10^4^ PFU) and treated with DNase I (10mg/kg, s.c) for 5 days. **(a)** Doublets, debris, and dead cells were first excluded. Leukocytes were identified as CD45+ events among viable cells. **(b)** Flow cytometry analyses of CD45+ living cells from the lung of K18-hACE2 infected mice treated or not with DNase I. Uninfected was used as control. Data are representative of two independent experiments and are shown as mean ± SEM. P values were determined by one-way ANOVA followed by Bonferroni’s post hoc test.

## REFERENCES

1. Wu Z, McGoogan JM. Characteristics of and Important Lessons From the Coronavirus Disease 2019 (COVID-19) Outbreak in China. JAMA [published online ahead of print: 2020]; doi:10.1001/jama.2020.2648

2. Guan W, et al. Clinical Characteristics of Coronavirus Disease 2019 in China. N. Engl. J. Med. [published online ahead of print: 2020]; doi:10.1056/nejmoa2002032

3. Dolhnikoff M, et al. Pathological evidence of pulmonary thrombotic phenomena in severe COVID-19. J. Thromb. Haemost. [published online ahead of print: 2020]; doi:10.1111/jth.14844

4. Batah SS, et al. COVID-19 bimodal clinical and pathological phenotypes. Clin. Transl. Med. 2022;12(1). doi:10.1002/CTM2.648

5. Skendros P, et al. Complement and tissue factor-enriched neutrophil extracellular traps are key drivers in COVID-19 immunothrombosis. J. Clin. Invest. [published online ahead of print: August 6, 2020]; doi:10.1172/JCI141374

6. Reusch N, et al. Neutrophils in COVID-19. Front. Immunol. 2021;12:952.

7. Schulte-Schrepping J, et al. Severe COVID-19 Is Marked by a Dysregulated Myeloid Cell Compartment. Cell 2020;182(6):1419–1440.e23.

8. Veras FP, et al. SARS-CoV-2–triggered neutrophil extracellular traps mediate COVID-19 pathologySARS-CoV-2 directly triggers ACE-dependent NETs. J. Exp. Med. 2020;217(12). doi:10.1084/jem.20201129

9. Middleton EA, et al. Neutrophil Extracellular Traps (NETs) Contribute to Immunothrombosis in COVID-19 Acute Respiratory Distress Syndrome.. Blood [published online ahead of print: June 29, 2020]; doi:10.1182/blood.2020007008

10. Zuo Y, et al. Neutrophil extracellular traps in COVID-19. JCI insight [published online ahead of print: 2020]; doi:10.1172/jci.insight.138999

11. Brinkmann V, et al. Neutrophil Extracellular Traps Kill Bacteria. Science (80-.). 2004;303(5663):1532–1535.

12. Jorch SK, Kubes P. An emerging role for neutrophil extracellular traps in noninfectious disease. Nat. Med. 2017;23(3):279–287.

13. Li P, et al. PAD4 is essential for antibacterial innate immunity mediated by neutrophil extracellular traps. J. Exp. Med. [published online ahead of print: 2010]; doi:10.1084/jem.20100239

14. Paryzhak S, et al. Neutrophil-released enzymes can influence composition of circulating immune complexes in multiple sclerosis. Autoimmunity [published online ahead of print: 2018]; doi:10.1080/08916934.2018.1514390

15. Saffarzadeh M, et al. Neutrophil extracellular traps directly induce epithelial and endothelial cell death: A predominant role of histones. PLoS One [published online ahead of print: 2012]; doi:10.1371/journal.pone.0032366

16. Henry CM, et al. Neutrophil-Derived Proteases Escalate Inflammation through Activation of IL-36 Family Cytokines. Cell Rep. 2016;14(4):708–722.

17. Toussaint M, et al. Host DNA released by NETosis promotes rhinovirus-induced type-2 allergic asthma exacerbation. Nat. Med. 2017;23(6):681–691.

18. Venkatesan P. Repurposing drugs for treatment of COVID-19. Lancet Respir. Med. 2021;9(7):e63.

19. MacConnachie AM. Dornase-alfa (DNase, Pulmozyme) for cystic fibrosis. Intensive Crit. care Nurs. 1998;14(2):101–102.

20. Ranasinha C, et al. Efficacy and safety of short-term administration of aerosolised recombinant human DNase I in adults with stable stage cystic fibrosis. Lancet [published online ahead of print: 1993]; doi:10.1016/0140-6736(93)92297-7

21. Holliday ZM, et al. Non-Randomized Trial of Dornase Alfa for Acute Respiratory Distress Syndrome Secondary to Covid-19. Front. Immunol. 2021;12. doi:10.3389/FIMMU.2021.714833

22. Dong W, et al. The K18-Human ACE2 Transgenic Mouse Model Recapitulates Non-severe and Severe COVID-19 in Response to an Infectious Dose of the SARS-CoV-2 Virus. J. Virol. 2022;96(1). doi:10.1128/JVI.00964-21

23. Golden JW, et al. Human angiotensin-converting enzyme 2 transgenic mice infected with SARS-CoV-2 develop severe and fatal respiratory disease. JCI Insight 2020;5(19). doi:10.1172/JCI.INSIGHT.142032

24. Zuo Y, et al. Neutrophil extracellular traps in COVID-19. JCI Insight [published online ahead of print: 2020]; doi:10.1172/jci.insight.138999

25. Zhang H, et al. Histopathologic Changes and SARS-CoV-2 Immunostaining in the Lung of a Patient With COVID-19. Ann. Intern. Med. [published online ahead of print: 2020]; doi:10.7326/M20-0533

26. Becker K, et al. Vasculitis and Neutrophil Extracellular Traps in Lungs of Golden Syrian Hamsters With SARS-CoV-2. Front. Immunol. 2021;12:1125.

27. Tian S, et al. Pathological study of the 2019 novel coronavirus disease (COVID-19) through postmortem core biopsies. Mod. Pathol. [published online ahead of print: 2020]; doi:10.1038/s41379-020-0536-x

28. Chen T, et al. Clinical characteristics of 113 deceased patients with coronavirus disease 2019: Retrospective study. BMJ [published online ahead of print: 2020]; doi:10.1136/bmj.m1091

29. Su H, et al. Renal histopathological analysis of 26 postmortem findings of patients with COVID-19 in China. Kidney Int. [published online ahead of print: 2020]; doi:10.1016/j.kint.2020.04.003

30. Hin Chu P†, et al. Comparative tropism, replication kinetics, and cell damage profiling of SARS-CoV-2 and SARS-CoV with implications for clinical manifestations, transmissibility, and laboratory studies of COVID-19: an observational study. The Lancet Microbe 2020;

31. Barnes BJ, et al. Targeting potential drivers of COVID-19: Neutrophil extracellular traps. J. Exp. Med. [published online ahead of print: 2020]; doi:10.1084/jem.20200652

32. Sanders JM, et al. Pharmacologic Treatments for Coronavirus Disease 2019 (COVID-19): A Review. JAMA 2020;323(18):1824–1836.

33. Fenton C, Keam SJ. Emerging small molecule antivirals may fit neatly into COVID-19 treatment. Drugs Ther. Perspect. Ration. drug Sel. use [published online ahead of print: February 28, 2022]; doi:10.1007/S40267-022-00897-8

34. Yang AP, et al. The diagnostic and predictive role of NLR, d-NLR and PLR in COVID-19 patients. Int. Immunopharmacol. [published online ahead of print: 2020]; doi:10.1016/j.intimp.2020.106504

35. Schneider AH, et al. Neutrophil extracellular traps mediate joint hyperalgesia induced by immune inflammation. Rheumatology (Oxford). 2021;60(7):3461–3473.

36. O’neil LJ, Kaplan MJ, Carmona-Rivera C. The Role of Neutrophils and Neutrophil Extracellular Traps in Vascular Damage in Systemic Lupus Erythematosus. J. Clin. Med. 2019;8(9). doi:10.3390/JCM8091325

37. Wong SL, et al. Diabetes primes neutrophils to undergo NETosis, which impairs wound healing. Nat. Med. [published online ahead of print: 2015]; doi:10.1038/nm.3887

38. Silva CM, et al. Gasdermin D inhibition prevents multiple organ dysfunction during sepsis by blocking NET formation. Blood [published online ahead of print: August 18, 2021]; doi:10.1182/BLOOD.2021011525/476604/GASDERMIN-D-INHIBITION-PREVENTS-MULTIPLE-ORGAN

39. Colón DF, et al. Neutrophil extracellular traps (NETs) exacerbate severity of infant sepsis. Crit. Care [published online ahead of print: 2019]; doi:10.1186/s13054-019-2407-8

40. Czaikoski PG, et al. Neutrophil extracellular traps induce organ damage during experimental and clinical sepsis. PLoS One [published online ahead of print: 2016]; doi:10.1371/journal.pone.0148142

41. Saitoh T, et al. Neutrophil extracellular traps mediate a host defense response to human immunodeficiency virus-1.. Cell Host Microbe [published online ahead of print: 2012]; doi:10.1016/j.chom.2012.05.015

42. Funchal GA, et al. Respiratory syncytial virus fusion protein promotes TLR-4-dependent neutrophil extracellular trap formation by human neutrophils. PLoS One [published online ahead of print: 2015]; doi:10.1371/journal.pone.0124082

43. Hiroki CH, et al. Neutrophil Extracellular Traps Effectively Control Acute Chikungunya Virus Infection. Front. Immunol. [published online ahead of print: 2020]; doi:10.3389/fimmu.2019.03108

44. Nie M, et al. Neutrophil Extracellular Traps Induced by IL8 Promote Diffuse Large B-cell Lymphoma Progression via the TLR9 Signaling. Clin. Cancer Res. 2019;25(6):1867–1879.

45. Keshari RS, et al. Cytokines Induced Neutrophil Extracellular Traps Formation: Implication for the Inflammatory Disease Condition. PLoS One [published online ahead of print: 2012]; doi:10.1371/journal.pone.0048111

46. Yipp BG, et al. Dynamic NETosis is Carried Out by Live Neutrophils in Human and Mouse Bacterial Abscesses and During Severe Gram-Positive Infection. Nat. Med. [published online ahead of print: 2015]; doi:10.1038/nm.2847

47. Hsia CCW, et al. An Official Research Policy Statement of the American ThoracicSociety/European Respiratory Society: Standards for Quantitative Assessment of LungStructure. Am. J. Respir. Crit. Care Med. 2010;181(4):394.

